# Microbe-dependent inter-organ communication regulates germline stem cell proliferation in *Drosophila*

**DOI:** 10.64898/2026.04.27.717131

**Authors:** Ritsuko Suyama, Miaomiao Zhao, Hiroshi Mori, Akiyoshi Hirayama, Shinichi Kawaguchi, Rizky M Fatimah, Yutaka Suzuki, Kazuharu Arakawa, Joanne Y Yew, Toshie Kai

**Affiliations:** Laboratory of Germline Biology, Graduate School of Frontier Biosciences, Osaka University, 1-3 Yamadaoka Suita, Osaka 565-0871, Japan; Advanced Genomics Center, National Institute of Genetics, 1111 Yata, Mishima, Shizuoka 411-8540, Japan; Institute for Advanced Biosciences, Keio University, Tsuruoka, Yamagata, 997-0017, Japan; Graduate School of Media and Governance, Keio University, Fujisawa, Kanagawa, 252-0882, Japan; Faculty of Environment and Information Studies, Keio University, Fujisawa, Kanagawa, 252-0882, Japan; Laboratory of Systems Genomics, Department of Computational Biology and Medical Sciences, Graduate School of Frontier Sciences, The University of Tokyo. 5-1-5 Kashiwanoha, Kashiwa, Chiba 277-8562,Japan; Pacific Biosciences Research Center, University of Hawai’i at Manoa, 1993 East-West Road, Honolulu, HI 96822, USA; Laboratory of Cell Recognition and Pattern Formation, Graduate School of Biostudies, Kyoto University, Sakyo-ku, Kyoto 606-8501, Japan

**Keywords:** germline stem cell, microbes, inter-organ communication, metabolism, circadian regulation, *Drosophila*

## Abstract

Animals integrate environmental cues with internal physiological states through inter-organ communication to regulate reproduction. However, how environmental microbes are incorporated into these systemic pathways to control reproductive stem cell proliferation remains poorly understood. Here, we demonstrate that environmental microbes colonizing the gut promote stem cell-mediated oogenesis in *Drosophila melanogaster* by increasing germline stem cell (GSC) number. This process requires microbial activation of gut metabolic pathways, including glycolysis and the pentose phosphate pathway (PPP), linking microbial cues to organismal physiology. We further show that microbial cues modulate circadian clock gene expression in both the gut and brain, and that circadian gene activity in these tissues is required for microbe-induced increases in GSCs. These metabolic changes promote ecdysone and juvenile hormone signaling in ovarian somatic cells, and single-cell transcriptomic analyses further reveal cell type-specific metabolic and hormonal responses across germline and follicle cell populations. Together, our findings establish that microbe-dependent gut-mediated inter-organ communication integrates metabolism, circadian gene expression, and endocrine signaling to regulate stem cell-mediated reproduction in *Drosophila*.

## Introduction

Germline stem cells (GSCs) are essential to *Drosophila* oogenesis, supplying the germ cells required for egg production and female fertility. The *Drosophila* ovary is among the most extensively characterized systems for dissecting stem cell regulation, providing fundamental insights into how specialized somatic niches coordinate stem cell maintenance, proliferation, and differentiation. GSCs reside at the anterior tip of the germarium, where somatic niche cells—including terminal filament cells (TF), cap cells (CC), and escort cells (EC)—support GSC self-renewal and regulate the balance between stem cell maintenance and differentiation^1^. Following asymmetric division, GSCs produce cystoblasts that undergo four rounds of incomplete cytokinesis to form 16-cell cysts, which develop into one oocyte and 15 nurse cells^1^. These cysts are encapsulated by follicle cells, and the oocyte matures through the transfer of mRNAs, proteins, and organelles from nurse cells^2,3^.

GSC activity is finely tuned by systemic physiological cues. Nutrient-rich diets accelerate oogenesis and enhance GSC proliferation, whereas nutrient-poor conditions impair these processes^4–10^. These effects are mediated largely by insulin-like peptides (Dilps) produced from neuroendocrine cells in the brain and gut, which promote germline and follicle-cell proliferation^4,6^. Mating provides an additional systemic trigger through neuroendocrine activation and ecdysone signaling to stimulate oogenesis^11–13^. The microbiome also shapes reproductive physiology by influencing nutrient intake, immunity, and metabolic homeostasis^14–20^. We previously demonstrated that environmentally acquired microbes enhance GSC proliferation and egg maturation via ecdysone- and juvenile hormone (JH)-dependent pathways, independently of insulin signaling^21^. Hormonal receptors for these pathways are expressed in germ cells and follicle cells, and their loss in the follicle cells impairs oogenesis^21–25^. Notably, microbe-induced GSC activation operates through somatic tissues and acts non–cell autonomously on GSC proliferation^21^. Thus, the *Drosophila* ovary provides a powerful model for dissecting how local somatic signals integrate with systemic cues to regulate stem cell behavior. These findings underscore the role of environmental cues in modulating GSC activity indirectly through inter-organ communication rather than by acting directly on the germline.

Inter-organ communication is a central mechanism through which environmental and physiological signals regulate reproductive function. In males, signals from the adjacent testis remodel intestinal carbohydrate metabolism via JAK–STAT signaling, generating a male-specific metabolic state that supports sperm development^26^. In females, dietary sugars modulate GSC proliferation through gut–germline communication. In addition, mating can trigger GSC proliferation through ovarian ecdysteroid production via the male-derived sex peptide^12^, while dietary glucose is converted to fructose in the hemolymph, which stimulates neuropeptide F (NPF) release and enhances GSC proliferation^27^. Together, these findings illustrate that metabolic and neuroendocrine signals originating outside the reproductive tissues profoundly influence GSC behavior via inter-organ pathways.

Microbes colonizing the gut can reshape host physiology by modulating energy balance, redox state, and biosynthetic capacity through diverse metabolites^28,29^. As a consequence, gut microbes influence nutrient utilization, immunity, and reproductive capacity across species^30–32^. In mammals, perturbation of the paternal gut microbiota compromises sperm quality and placental development. These defects are reversed by restoring the microbiota prior to conception, thereby rescuing fetal development^33^. Microbe–brain communication is also well documented, as gut microbes alter neural activity, behavior, and mating preferences through metabolic and neuroendocrine pathways^34–36^. Together, these studies indicate that environmental microbes can function as regulators of inter-organ communication and are capable of modulating systemic metabolic and reproductive physiology.

In parallel, host metabolic state and circadian timing are tightly interconnected. Core metabolic pathways such as glycolysis and gluconeogenesis exhibit circadian regulation^37^, while metabolic outputs, including NAD⁺ levels and cellular redox status, feed back to the core clock machinery^38–43^. Disruption of this metabolic–temporal coupling can lead to physiological decline and metabolic dysfunction^40,44–47^. Taken together, these observations suggest that interactions with circadian clock machinery may represent an additional regulatory layer through which environmental microbes modulate host metabolic physiology. Despite extensive evidence that gut microbes regulate multiple host organ systems, the molecular mechanisms linking microbial cues from the gut and brain to ovarian function remain unclear. In particular, how these systemic signals are integrated to regulate reproductive stem cell behavior has yet to be defined.

Here, we demonstrate that environmentally acquired microbes promote oogenesis by modulating inter-organ communication linking the gut, brain, and ovary in *Drosophila melanogaster*. Bulk transcriptomic analyses revealed that microbial exposure activates glycolysis, the pentose phosphate pathway (PPP), and circadian programs in the gut and brain, which in turn modulate ecdysone and juvenile hormone (JH) signaling―key regulators of GSC homeostasis. Functional experiments further showed that gut-derived glycolytic and PPP activities, together with circadian clock regulators in both gut and brain, are required for microbe-induced GSC proliferation. Single-cell transcriptomic profiling of the ovary uncovered stage- and cell type–specific transcriptional responses to microbial cues across germline and somatic lineages. Together, these findings define a microbe–gut–brain–ovary axis in which metabolic and circadian gene regulation converge to control reproductive stem cell activity.

## Results

### Microbial cues induce tissue-specific transcriptional programs in the gut and brain

We previously demonstrated that environmental microbes promote germline stem cell (GSC) proliferation and accelerate oogenesis in *Drosophila*^21^. To elucidate the molecular basis underlying this response, we employed a previously established exposure scheme in which donor flies were maintained in vials for 3–4 days, after which newly eclosed virgin females (“Acceptor”) were replaced into these vials for three days (Fig. 1A). Consistent with our prior observations, acceptor females exposed to microbe-rich (M+) vials exhibited a significant increase in GSC number compared with those maintained under microbe-poor (M-) conditions^21^. To identify tissues responsive to microbial cues, we performed bulk RNA sequencing on dissected guts, brains, and ovaries from M+ and M– females (three biological replicates per condition; Fig. 1B). After quality control and alignment to the *Drosophila melanogaster* reference genome (BDGP ver. 6.32), differential gene expression analyses were performed independently for each tissue.

**Figure. 1.**
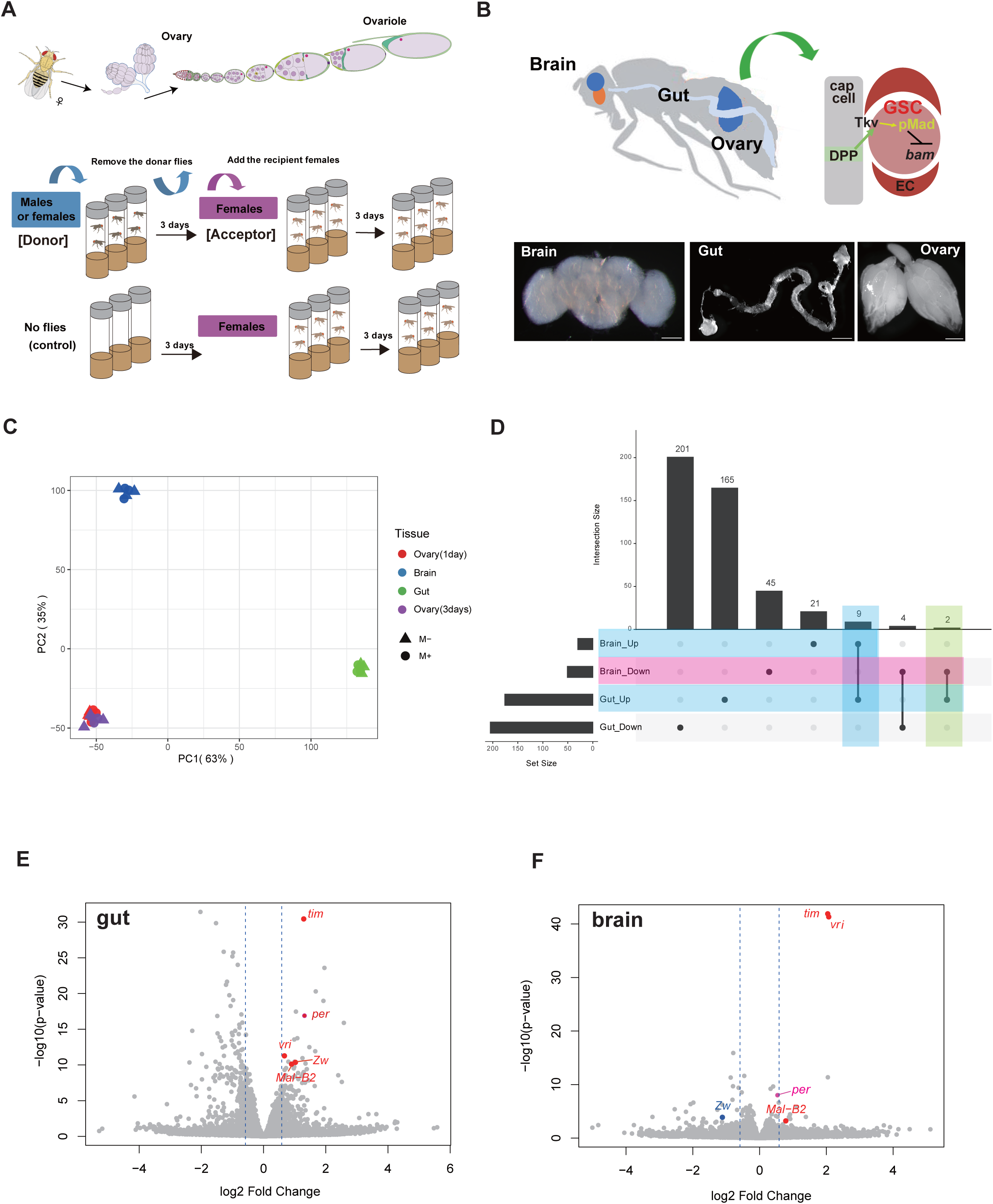

Principal component analysis (PCA) of the top 500 differentially expressed genes revealed clear clustering of samples by tissue type (Fig. 1C), with the strongest segregation between M+ and M- conditions observed in the gut (Supplementary Fig. 1A). Heatmap analysis confirmed high reproducibility among biological replicates within each tissue (Supplementary Fig. 1B). Differential expression analysis (log2 fold change ≥1.5, *p* < 0.01) showed that the gut exhibited the most pronounced transcriptional response to microbial exposure, with 201 genes upregulated and 165 genes downregulated under M+ conditions compared with M- conditions (Fig. 1D). The brain also showed substantial transcriptional changes, with 21 genes upregulated and 45 downregulated. In contrast, the ovary exhibited relatively few differentially expressed genes, with limited overlap across exposure durations. Comparing M+ to M− conditions, 18 genes were upregulated and 19 were downregulated in ovaries after one day of exposure, whereas two genes were upregulated and 93 were downregulated after three days (Supplementary Fig. 1C).

Because transcriptional differences in the ovary were modest and variable between one- and three-day exposures (Supplementary Fig. 1B-E), we focused subsequent bulk transcriptomic analyses on the gut and brain. Comparison of these two tissues identified a small subset of genes that were consistently regulated under M+ conditions. Nine genes were commonly upregulated and four were commonly downregulated in both the gut and brain (Fig. 1D). Notably, among the shared upregulated genes were circadian regulators (*vri*, *tim*) and a glycolytic and pentose phosphate pathway (PPP)-associated gene (*Mal-B2*), whereas *zw* was upregulated in the gut but downregulated in the brain (Fig. 1E,F; Supplementary Table 1). These coordinated transcriptional changes were not readily detected in ovarian bulk RNA-seq, likely reflecting the cellular heterogeneity and asynchronous developmental states within ovarian tissue rather than absence of microbial effects (Supplementary Fig. 1D,E). As ovarian bulk RNA-seq showed only modest transcriptional changes, we used single-cell RNA sequencing in subsequent analyses to resolve cell type– and stage-specific ovarian responses to microbial cues (see below).

### Gut metabolic pathways are required for microbe-induced GSC proliferation

Bulk RNA-seq analyses identified PPP and glycolysis as candidate metabolic pathways responsive to microbial exposure in the gut. To determine whether these metabolic pathways are functionally required for microbial stimulation of GSC proliferation, we quantified GSC numbers following tissue-specific knockdown of PPP- and glycolysis-associated genes. Using the enteroendocrine cell (EE)-specific driver *TKg-Gal4,* which is expressed in the gut^11,48,49^, knockdown of key PPP enzymes (*Mal-B2, hex-A, zw, 6PGL,* and *pgd*) significantly suppressed the microbe-induced increase in GSC number (Fig. 2A). Similarly, knockdown of glycolytic genes (*pgi, pfk,* and *pyk*) reduced GSC numbers under M+ conditions (Fig. 2B), indicating that both metabolic pathways are required in EEs for microbe-induced GSC proliferation. We next examined whether these metabolic pathways are required in another gut cell population. Using *mex-1-Gal4*, which is expressed in enterocytes^48,49^, knockdown of PPP or glycolytic enzymes likewise abolished the microbe-dependent increase in GSC number (Fig. 2C,D). These results indicate that microbial regulation of gut metabolism in both EEs and enterocytes is essential for promoting GSC proliferation in the ovary.

**Figure. 2.**
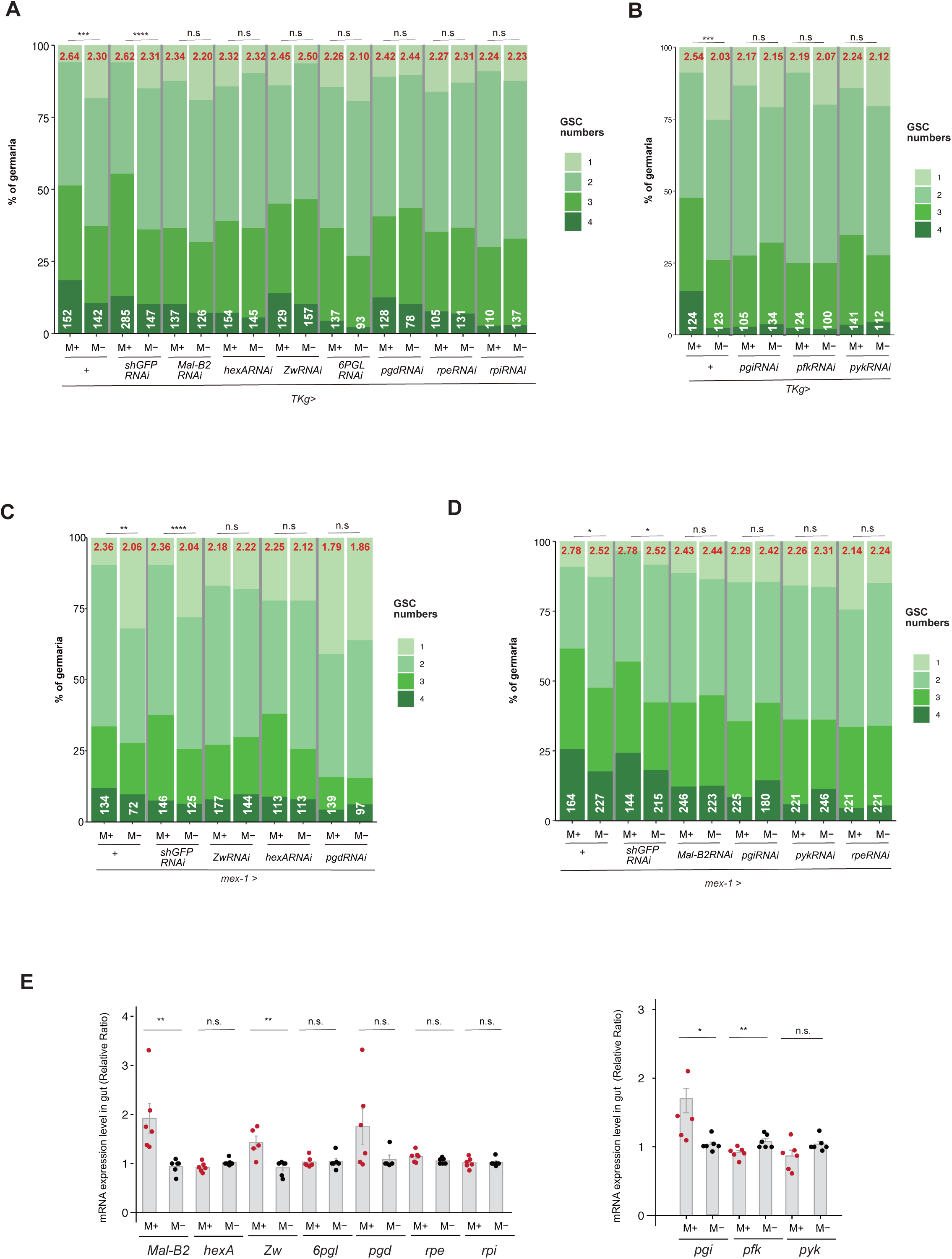

Previous studies reported that PPP activity is required for late oogenesis and egg production, whereas glycolysis appears largely dispensable at these stages^50^. To determine whether PPP activity also contributes to early germline regulation, we attenuated PPP gene expression in germ cells using *Nanos* (*NN*)-Gal4–driven RNA interference. Knockdown of *Mal-B2*, *Hex-A*, or *Zw* significantly suppressed the GSC increase normally observed under M+ conditions, reducing GSC numbers to levels comparable to M– controls (Supplementary Fig. 2A). These findings indicate that PPP activity is required for late oogenesis and egg maturation, and that early germline proliferation depends on PPP function both within the ovary and through gut-derived signals. To validate transcriptional changes in these pathways, we quantified representative PPP and glycolytic genes by qPCR in guts from M+ and M– females. Consistent with the RNA-seq data, *Mal-B2* and *Zw* were significantly upregulated in the gut under M+ conditions, and *Pgi* showed a modest increase (Fig. 2E). Other genes in these pathways showed minimal changes, consistent with the RNA-seq profiles. Nevertheless, PPP and glycolytic genes expressed in the gut were functionally required for the microbe-induced activation of GSC proliferation. Together, these results indicate that microbial stimulation promotes early oogenesis via gut-derived metabolic signaling together with germline-autonomous metabolic activity.

### Microbes reshape intestinal stem cell differentiation and gut endocrine potential

To determine where microbes localize after exposure to a microbe-rich environment, we quantified microbial titers in dissected guts and ovaries (Supplementary Fig. 3A). In both male and female non-mated donor flies, microbes were predominantly detected in the gut and were undetectable in the ovaries. A similar distribution was observed in recipient females maintained under M+ and M− conditions: microbial load was consistently confined to the gut and was markedly elevated under M+ conditions. No microbial signal was detected in the ovary under any condition, indicating that microbial effects are mediated indirectly via extra-ovarian tissues rather than by direct ovarian colonization.

To assess how microbial exposure alters gut physiology, we examined intestinal stem cell (ISC) differentiation using established lineage markers and nuclear size criteria^51^. The *esg*-Gal4-driven *UAS-mCD8::GFP* (*esg>mCD8::GFP*) labels ISCs and progenitor populations, including enteroblasts (EBs) and pre–enteroendocrine cells (preEEs), whereas Prospero (Pros) marks preEEs and mature enteroendocrine cells (EEs). Accordingly, ISCs and EBs were GFP-positive (GFP⁺) but Pros-negative (Pros⁻), preEEs were GFP⁺/Pros⁺, EEs were GFP⁻/Pros⁺, and polyploid enterocytes were GFP⁻/Pros⁻ with large nuclei (Fig. 3A).

**Figure. 3.**
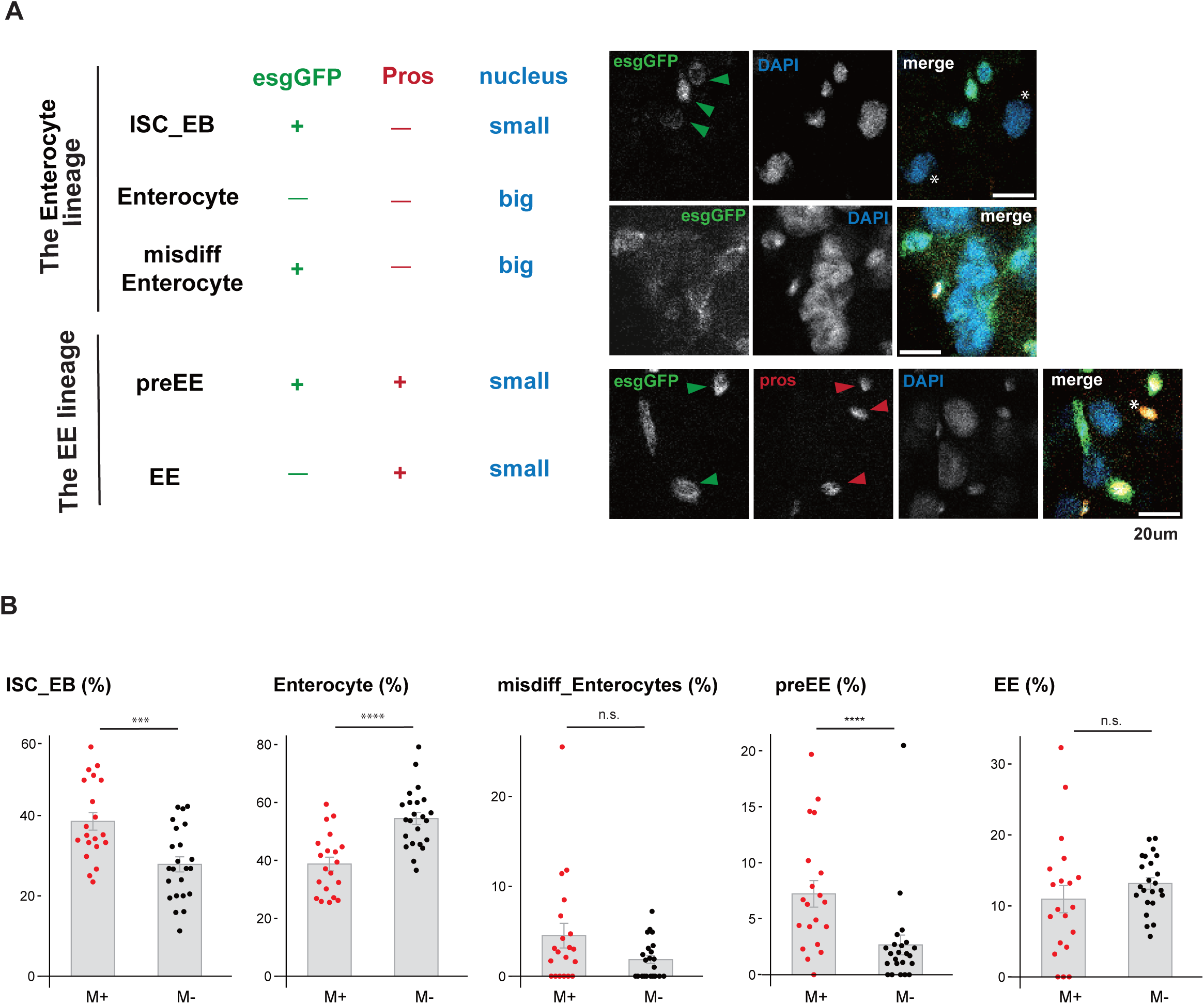

Microbial exposure altered specific branches of the ISC lineage (Fig. 3B). The proportion of ISCs/EBs increased by ∼40% under M+ conditions relative to M− controls (M+, 38.6 ± 9.6%; M−, 27.9 ± 8.8%; *P* < 0.005), and preEEs increased by ∼2.5-fold (M+, 7.2 ± 5.2%; M−, 2.7 ± 4.1%; *P* < 0.005). In contrast, the proportion of mature EEs did not differ significantly between conditions. Notably, the fraction of enterocytes was reduced by ∼30% in the M+ gut (M+, 38.8 ± 10.3%; M−, 54.5 ± 10.0%; *P* < 0.001). Together, these results demonstrate that microbial exposure reshapes ISC differentiation, favoring early EE-lineage progenitors while reducing the relative abundance of mature enterocytes. This shift in the proportions of intestinal cell types suggests that microbial cues remodel gut physiological output, potentially influencing endocrine and metabolic signaling to distal organs.

### Microbial exposure enhances PPP-linked metabolic outputs in the gut

To determine whether transcriptional changes in gut PPP and glycolysis are reflected at the metabolic level, we performed comprehensive profiling of anionic and cationic metabolites in the gut, where microbes localize and intestinal cell differentiation is altered under microbe-rich conditions. Principal component analysis (PCA) revealed clear separation between gut samples from M+ and M− females (four biological replicates per condition), indicating clear microbe-dependent metabolic remodeling (Supplementary Fig. 3B,C). Despite substantial variability among individual metabolites, central carbon metabolism displayed a coherent and reproducible shift under M+ conditions. Notably, PPP-associated metabolites showed a selective increase, whereas glycolysis and the tricarboxylic acid (TCA) cycle showed an overall reduction (Supplementary Fig. 4A–E). Consistent with increased PPP flux, the PPP intermediate sedoheptulose-7-phosphate (S7P) and the major PPP output NADPH were significantly elevated in M+ guts (Supplementary Fig. 4A,B,F, Supplementary Fig. 5A,D).

In contrast, NADH levels and several glycolytic intermediates tended to be reduced in M+ guts, although these changes did not reach statistical significance (Supplementary Fig. 4C,F, Supplementary Fig. 5D). While a large fraction of both anionic and cationic metabolites showed an overall downward trend under M+ conditions (Supplementary Fig. 3C, Supplementary Fig. 4A,B,F–H, Supplementary Fig. 5A–D), this global decrease contrasted with the selective upregulation of PPP-associated metabolites. Of note, several amino acid- and polyamine-related metabolites, including kynurenine and Gly-Gly, were increased under M+ conditions (Supplementary Data 3), indicating that specific metabolic pathways are affected by microbial activity rather than a uniformly suppressed. In addition, riboflavin consistently exhibited an upward trend in M+ guts, mirroring its behavior in ovaries (Supplementary Fig. 4F, Supplementary Fig. 5D). This observation is consistent with previous reports that microbial exposure enhances mitochondrial function to support ATP production and oogenesis^20,21^.

### Circadian genes in the gut and brain are required for microbe-induced GSC proliferation

We next examined whether circadian clock gene activity provides an additional regulatory layer that modulates the metabolic–endocrine axis in response to microbial exposure. Bulk RNA-seq analysis revealed that multiple core circadian clock genes were upregulated in both the gut and brain under microbe-rich conditions (Fig. 1E,F; Supplementary Table 1). To assess the functional relevance of this activation, we disrupted circadian gene expression in these tissues and quantified GSC numbers. Knockout or knockdown of core circadian regulators (*per, tim, or vri*) in enterocytes using *mex-1-Gal4* abolished the microbe-induced increase in GSC number (Fig. 4A). Similarly, perturbation of these genes in the gut and brain using *tim-Gal4* significantly suppressed GSC increase under M+ conditions (Fig. 4B). In contrast, control manipulations targeting *acp* or *shGFP* did not affect GSC number under M+ conditions with either driver. In addition, no suppression of GSC numbers was observed following disruption of core circadian regulators using *TKg-Gal4,* consistent with the lack of expression of these genes in the EE cells (Supplementary Fig. 2B). These results indicate that circadian gene activity in gut enterocytes is specifically required for microbe-dependent GSC proliferation.

**Figure. 4.**
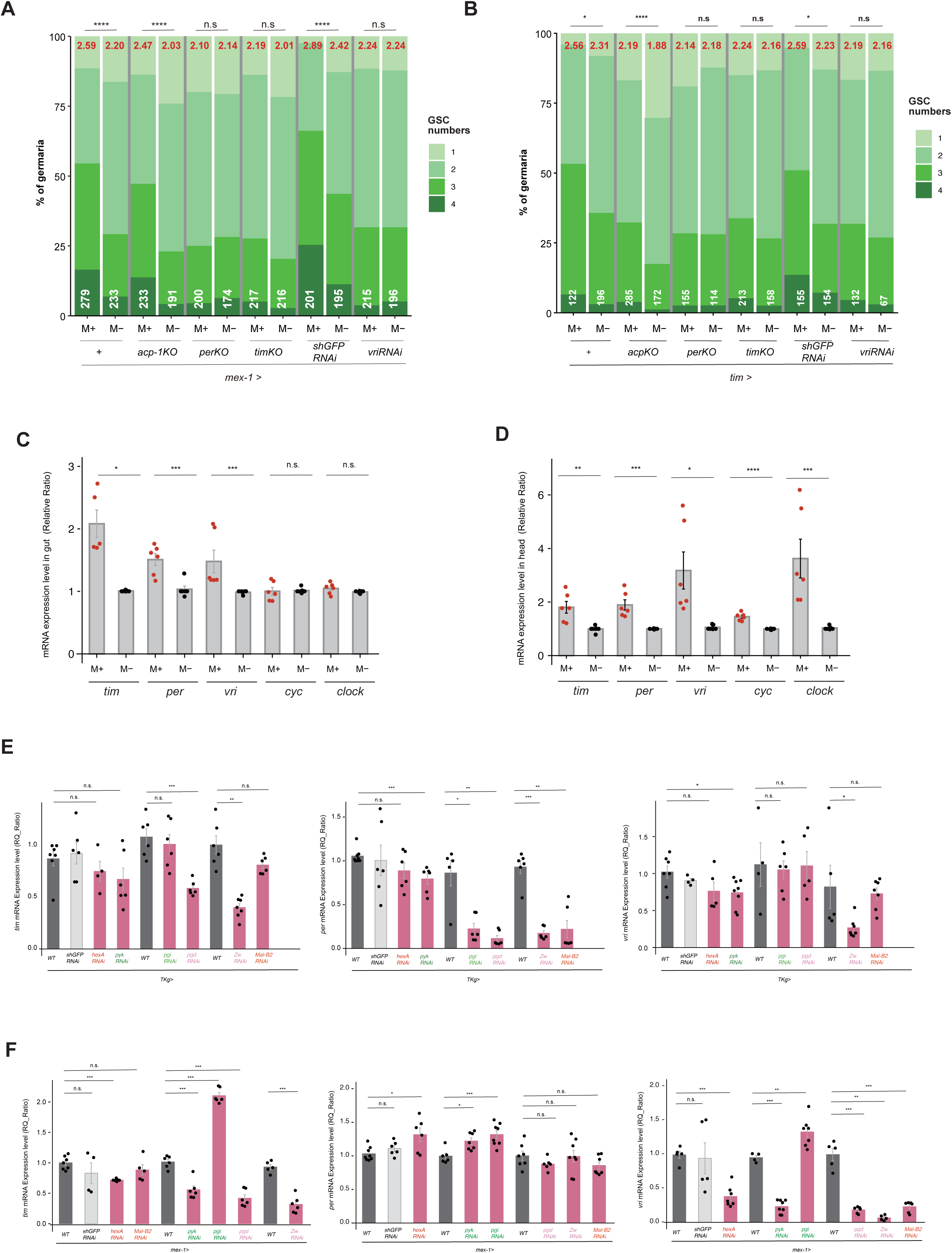

Because circadian programs can be differentially regulated across tissues and are not necessarily synchronized in phase or amplitude^52–54^, we next examined whether microbial exposure alters circadian gene expression in the gut and brain. Quantitative analysis revealed that *tim*, *per*, and *vri* transcripts were significantly upregulated in the gut of M+ females, whereas other clock components (*cyc*, *clock*) remained unchanged (Fig. 4C). A similar upregulation of *tim*, *per*, and *vri* was observed in heads from the same animals, whereas *cyc* and *clock* were also upregulated (Fig. 4D). These results confirm that microbial exposure enhances expression of circadian clock genes across gut and brain, consistent with the RNA-seq results.

Because metabolic pathways such as PPP and glycolysis can influence circadian oscillations through redox-sensitive mechanisms^39,42,43,55,56^, we next asked whether gut-derived metabolic activity contributes to circadian clock gene regulation in the brain. Knockdown of PPP- or glycolysis-associated genes in enteroendocrine cells (*TKg-Gal4*) or enterocytes (*mex-1-Gal4*) significantly reduced the expression of several circadian genes in the head (Fig. 4E,F). Notably, *TKg-Gal4***–**mediated perturbation preferentially reduced *per* expression, whereas *mex-1-Gal4*–driven knockdown reduced *tim* and *vri* expression more strongly, suggesting that distinct gut cell types contribute differently to gut–brain circadian signaling. Together, these results demonstrate that microbial activation of circadian rhythm programs in the gut is required for GSC proliferation and that gut metabolic activity contributes to circadian clock gene regulation in the brain. These findings suggest a functional link between peripheral metabolic pathways, regulation of circadian clock genes, and microbe-induced control of germline stem cell activity.

### Gut-derived metabolic signaling regulates ovarian ecdysone and JH receptor activity

Because the microbe-induced increase in GSCs was abolished by functional downregulation of PPP and glycolysis pathways in the gut (Fig. 2A–D), we next examined how gut-derived metabolic signals influence ovarian physiology. In our previous study, we showed that both ecdysone and juvenile hormone (JH) signaling pathways are activated in ovaries of females maintained under microbe-rich conditions, using receptor-conjugated *LacZ* reporters^21^. To test whether gut metabolic pathways are required for ovarian hormone receptor activation, we employed heat shock–inducible *ecR-LacZ* and *gce-LacZ* reporters, visualized using the fluorescent β-galactosidase substrate SPiDER-βGal^21,57,58^.

In females carrying *ecR-LacZ*, knockdown of *Zw* in either enteroendocrine cells (*TKg-Gal4*) or enterocytes (*mex-1-Gal4*) resulted in comparable *LacZ* signal intensities between M+ and M− conditions in anterior germarial somatic cells, thereby abolishing the enhancement of EcR reporter activity observed under M+ conditions (Fig. 5A). In contrast, wild-type females maintained under M+ conditions displayed robust EcR reporter activation, consistent with our previous observations (Supplementary Fig. 5E)^21^. A similar suppression of EcR activation was observed upon gut-specific knockdown of *Mal-B2* (Fig. 5B). These results indicate that microbial stimulation of ecdysone signaling in early ovarian somatic cells requires activation of metabolic pathways in the gut.

**Figure. 5.**
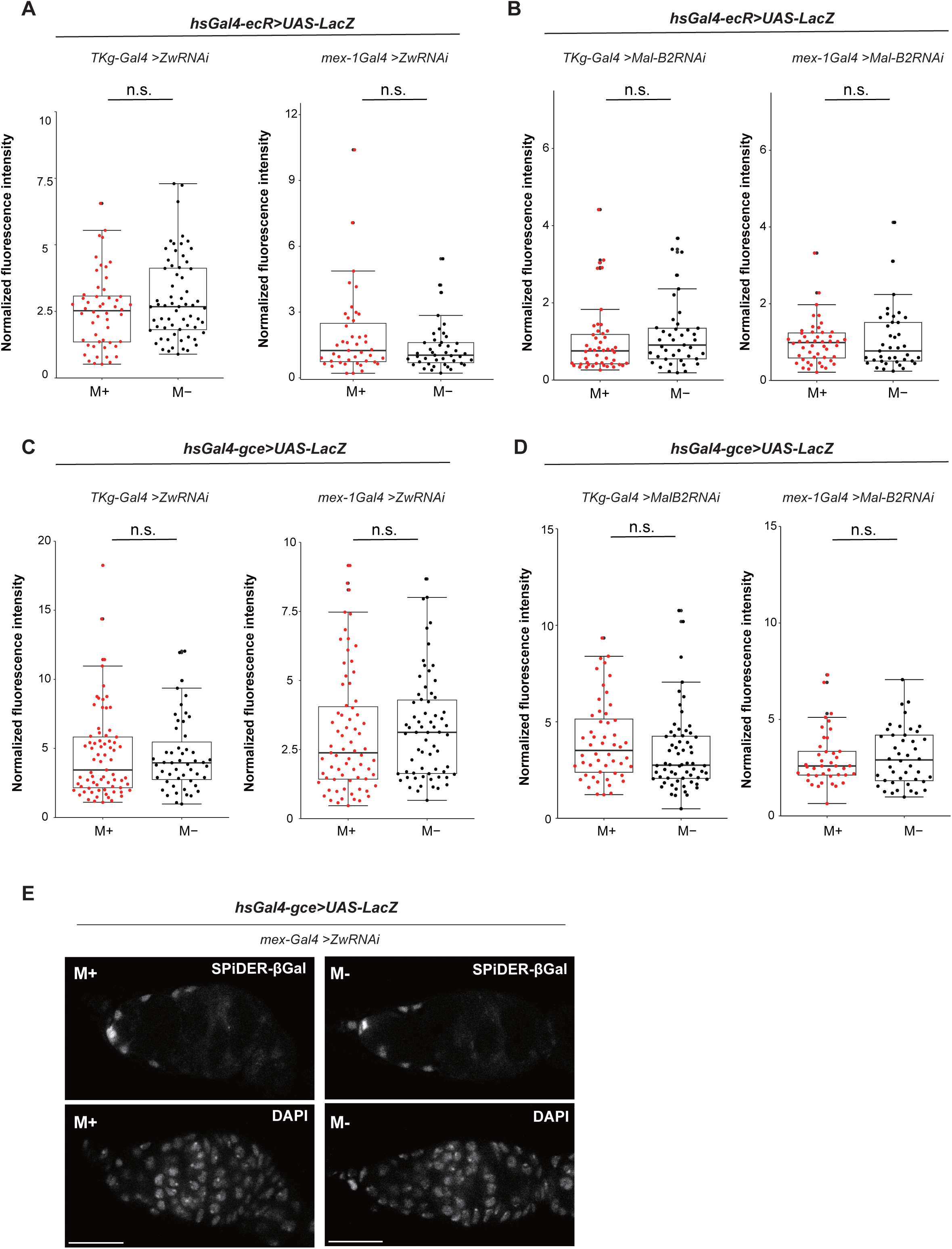

We next tested whether microbial cues similarly regulate JH signaling using the *gce-LacZ* reporter. As observed for EcR, JH receptor activity was markedly reduced under M+ conditions when *Zw* or *Mal-B2* was knocked down by either *TKg-Gal4* or *mex-1-Gal4,* as indicated by reduced *LacZ* signal intensity (Fig. 5C–E). In contrast, wild-type females maintained under M+ conditions exhibited strong activation of the *gce* reporter (Supplementary Fig. 5F). The concordant suppression of both EcR and JH receptor activation upon disruption of gut metabolic pathways demonstrates that gut-derived metabolic activity is necessary for ovarian hormonal signaling. Together, these findings support a model in which microbial activation of gut metabolism regulates both ecdysone and JH signaling in ovarian somatic cells.

### Single-cell transcriptomics identifies ovarian somatic and germline responses to microbial cues

Finally, to examine how microbial cues are reflected across individual ovarian cell types at single-cell resolution, we performed single-cell RNA sequencing (scRNA-seq) on ovaries from females maintained under M+ and M- conditions. Because bulk RNA-seq analysis revealed only modest and variable transcriptional changes in whole ovaries, likely reflecting cellular heterogeneity and asynchronous developmental states, we performed scRNA-seq to resolve cell type–specific responses to microbial exposure and clarify how inter-organ communication regulates oogenesis.

Early-stage ovarian tissues were dissociated and subjected to scRNA-seq (Fig. 6A). After quality control (see Methods), 7,600–8,000 cells from M+ samples and 5,000–7,000 cells from M− samples were retained for downstream analyses. Uniform Manifold Approximation and Projection (UMAP) embedding of the merged datasets revealed well-separated clusters corresponding to the major germline and somatic lineages of the *Drosophila* ovary (Fig. 6B)^59^. Cell-type identities were assigned based on established marker genes from previous ovarian single-cell studies (Fig. 6B,C,E-I; Supplementary Fig. 6A,B, see Methods) and were further confirmed by dot plots, feature plots, and violin plots (Fig. 6D,E-I; Supplementary Fig. 6A,B)^60–64^. Consistent with this annotation, microbe-dependent transcriptional changes were restricted to specific cell types and developmental stages, whereas most genes showed comparable expression between M+ and M-conditions. Importantly, the identified clusters were consistent with known developmental relationships within both germline and somatic lineages and captured continuous transcriptional progression across ovarian developmental stages. Together, these results establish a well-annotated single-cell atlas of early-stage ovaries under M+ and M- conditions, enabling identification of cell type– and stage-specific transcriptional responses to microbial cues.

**Figure. 6.**
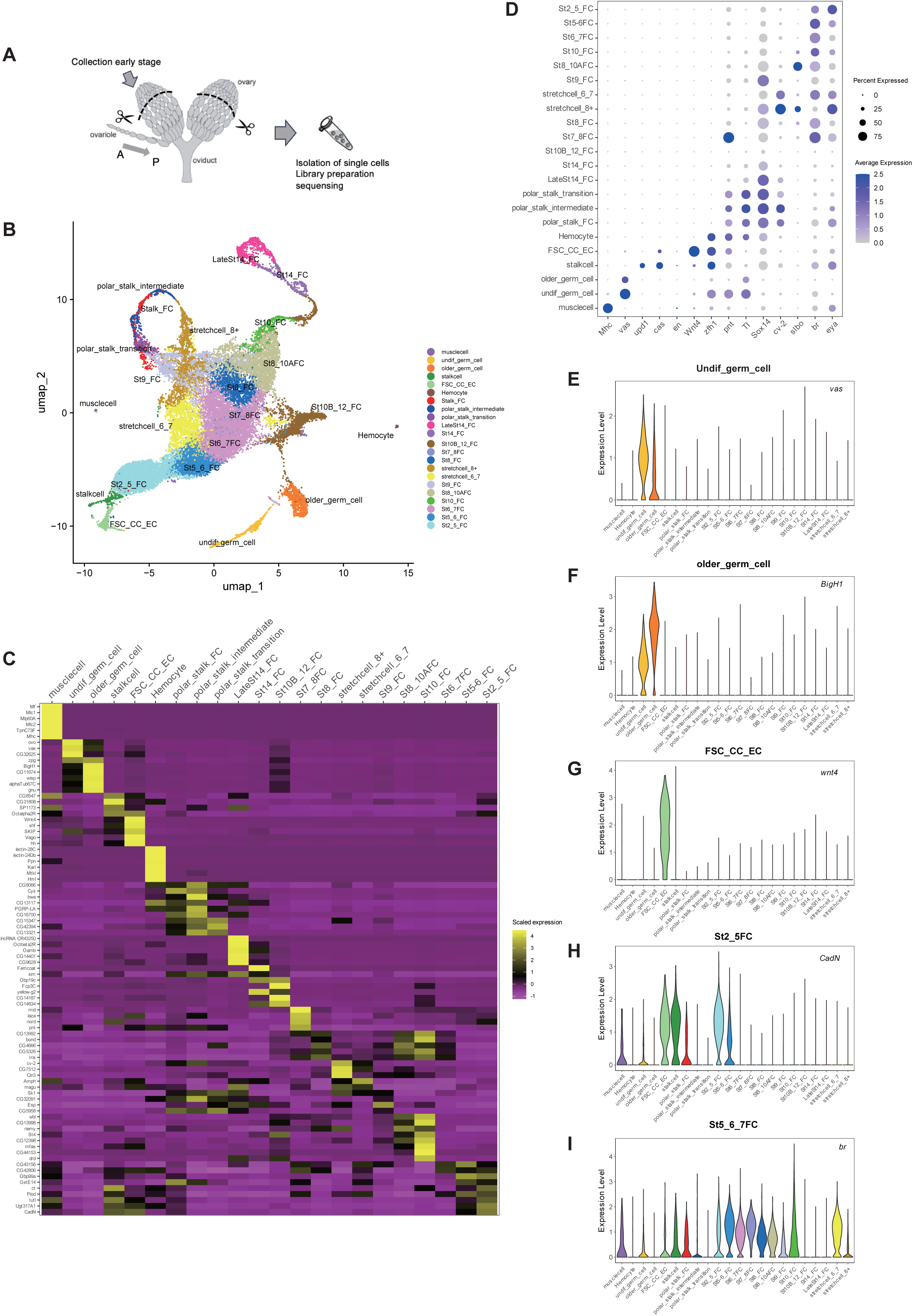

To examine how microbial exposure influences ovarian transcription at cellular resolution, we performed differential gene expression (DEG) analyses between M+ and M− conditions for each annotated cell type. Consistent with bulk RNA-seq results, microbial exposure did not induce uniform transcriptional changes across ovarian cell populations. Instead, microbe-dependent responses were restricted to specific cell types and developmental stages, with transcriptional changes more frequently observed in late somatic lineages (Supplementary Fig. 7A,B).

To refine germline-specific responses, we subclustered the germline lineage and identified three populations corresponding to germline stem cells (GSCs), undifferentiated germ cells, and older differentiated germ cells (Fig. 7A,B). Although GSCs exhibited relatively few DEGs, under M+ conditions, undifferentiated germ cells displayed significant upregulation of glycolytic genes, including *Eno*, with similar upward trends observed in GSCs and differentiated germ cells (Fig. 7C; Supplementary Fig. 7C). Pairwise DEG analyses further identified glycolysis-related genes, *Pyk* and *Pep,* as upregulated in undifferentiated germ cells, whereas *foxo*, a key component of insulin signaling, was downregulated (Fig. 7D-F). To capture coordinated pathway-level responses, we analyzed module scores for glycolysis, PPP, and hormone-related pathways across germline subtypes (Fig. 7G). Glycolytic activity was significantly increased in undifferentiated germ cells under M+ conditions (Δmedian = +0.094; Wilcoxon FDR = 0.015) and showed a weaker but detectable elevation in GSCs (Fig. 7G; Supplementary Fig. 7D). Hormone-response and germline-regulatory modules were selectively elevated in older germ cells (Hormone: FDR = 0.0035; Germline: FDR = 0.0106) (Fig. 7G; Supplementary Fig. 7D), whereas PPP activity exhibited only a modest increase in GSCs that did not reach significance after multiple-testing correction (Δmedian = +0.11, p = 0.037; FDR = 0.11)(Fig. 7G; Supplementary Fig. 7D). These results indicate that microbial cues elicit stage-specific metabolic and signaling responses within the germline rather than uniform transcriptional activation. In particular, glycolysis was preferentially enhanced in undifferentiated germ cells, whereas hormone and germline regulatory signatures were activated in older germ cells. PPP activity showed a modest shift in GSCs, consistent with metabolic and signaling heterogeneity across germline differentiation status.

**Figure. 7.**
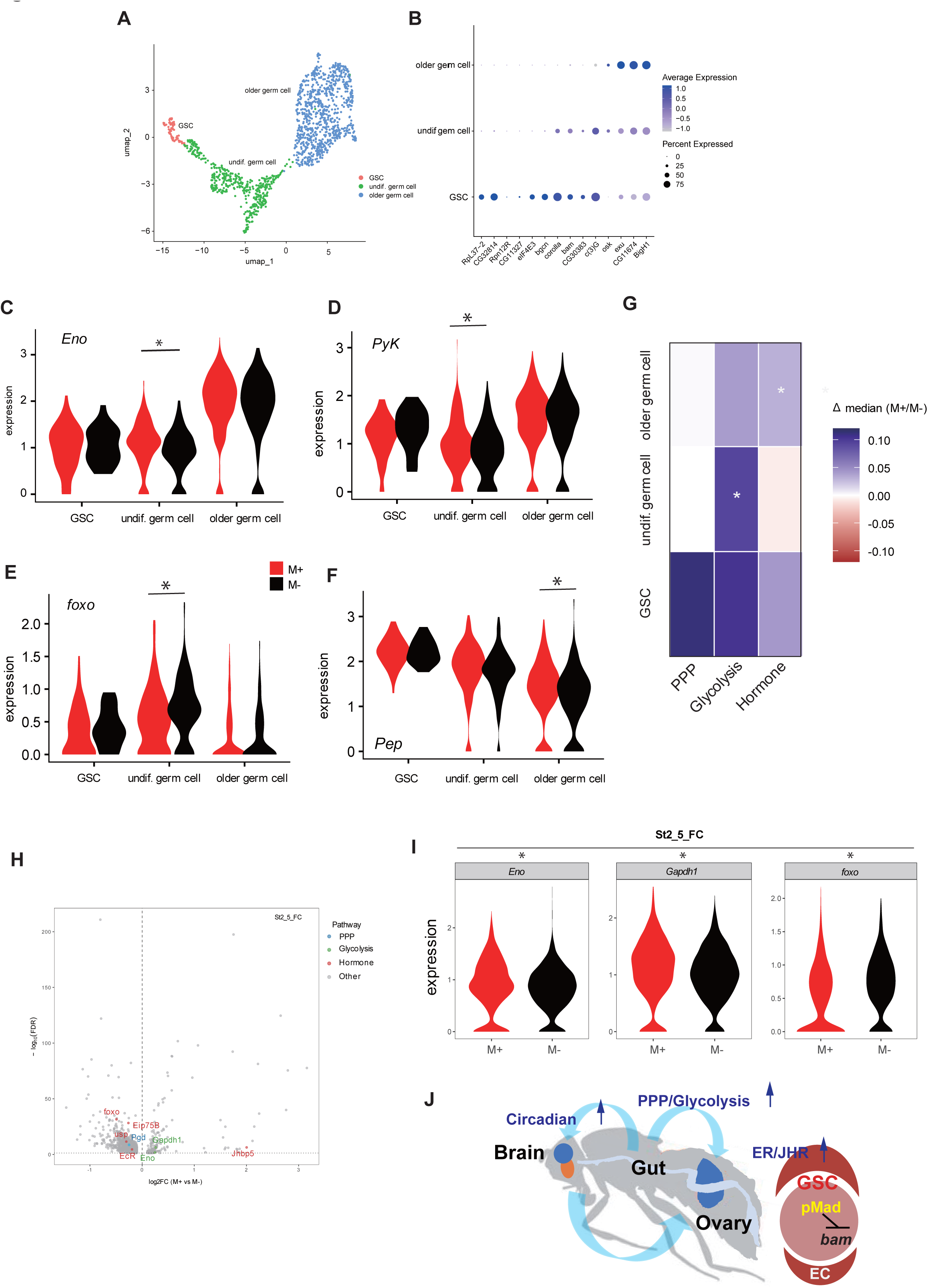

Among somatic populations, transcriptional changes were most prominent in early and mid-stage follicle cells, and were largely characterized by reduced expression of a subset of genes under M+ conditions (Supplementary Fig. 7A). Analysis of metabolic and hormone-associated genes revealed a tendency toward increased expression of glycolytic enzymes in somatic lineages, whereas the juvenile hormone-binding protein *Jhbp5* was elevated in both early germline cells and somatic follicle populations (Supplementary Fig. 7B), consistent with enhanced hormonal signaling under M+ conditions.

In contrast to germline populations, somatic niche cells identified after subclustering, including follicle stem cells (FSCs), terminal filament/cap cells (TF–CC), and ECs, showed few DEGs between conditions (Supplementary Fig. 7E,F). Despite this overall transcriptional stability, pathway-level analyses revealed subtle but reproducible shifts in metabolic and hormone-related signatures across somatic lineages, including ECs positioned adjacent to early germ cells. Specifically, PPP activity was modestly increased in TF–CC and ECs, while hormone-related signatures showed small yet consistent changes in FSCs and TF–CC (Supplementary Fig. 7G).

Consistent with these pathway-level changes, Gene Ontology enrichment analyses supported activation of metabolic and signaling processes in specific cell types. More pronounced microbe-dependent transcriptional changes were observed during follicle-cell differentiation, where mid-staged follicle cells (stage 2–5) exhibited elevated expression of glycolytic enzymes such as *Eno* and *Gapdh1*, accompanied by downregulation of *foxo*, indicative of enhanced glycolytic activity and attenuated insulin signaling (Fig. 7H,I; Supplementary Fig. 7H). Gene Ontology analysis of these stage 2–5 follicle cells revealed significant enrichment of nucleotide and energy metabolism–related biological processes (Supplementary Fig. 7I). These pathways are consistent with increased biosynthetic and energetic demand supported by glycolytic and PPP flux. Collectively, these results indicate that microbial cues modulate glycolytic, hormonal, and associated signaling responses in defined germline and somatic populations in a stage-dependent manner rather than globally altering ovarian gene expression.

## Discussion

Animals must integrate fluctuating environmental inputs into stable reproductive decisions to ensure species continuity. *Drosophila* acquire microbes from environmental sources such as dietary substrates, frass, and conspecific contact^15,65^. Here, we propose that environmental microbes promote *Drosophila* GSC proliferation through a microbe–gut–brain–ovary inter-organ communication network. Using integrated transcriptomics, genetic manipulation, metabolomics, and ovarian single-cell profiling, we demonstrate that microbial exposure remodels gut metabolism, coordinates gut–brain circadian clock regulation, and potentiates endocrine signaling in ovarian somatic cells to support GSC proliferation.

The gut functions as a metabolic sensor that translates microbial signals into systemic reproductive outputs which are coordinated with organismal metabolic state. In *Drosophila*, previous studies have shown that intrinsic activation of the PPP in the germline supports egg maturation and can influence organismal metabolism by modulating fat-body–derived satiety factors such as *fit*^50,66^. In contrast, our findings reveal that GSC proliferation is also influenced by extrinsic metabolic inputs originating in the gut, where microbial exposure activates glycolysis and PPP processes. Suppressing either pathway in enteroendocrine cells or enterocytes of the gut abolished the microbe-induced GSC increase, demonstrating that the gut transduces microbial signals to promote germline proliferation. These extrinsic metabolic inputs act alongside cell-autonomous metabolic requirements in the germline, together supporting GSC proliferation (Supplementary Fig. 2A, Supplementary Fig. 7C)^67^. Activation of these pathways may provide both energetic and biosynthetic substrates that facilitate endocrine signaling within the ovarian niche (Fig. 5).

Notably, steady-state levels of glycolytic intermediates were not elevated in M+ guts (Supplementary Fig. 4C-E), despite genetic and functional evidence showing that glycolytic activity is required for microbe-induced GSC proliferation (Fig. 2B,D). This discrepancy is consistent with the flux-controlled nature of glycolysis, in which pathway activity is not necessarily reflected by metabolite abundance^68–70^. Glycolytic flux can supply glucose-6-phosphate to the PPP, supporting NADPH production and biosynthetic capacity without global accumulation of glycolytic intermediates^71–74^. Consistent with this model, NADPH and the PPP intermediate S7P were selectively elevated under microbe-rich conditions, despite an overall reduction in many other metabolites (Supplementary Fig. 4A–F, Supplementary Fig. 5A,D). Together, these findings suggest that microbial cues regulate gut metabolism primarily through changes in metabolic flux rather than metabolite accumulation.

The mechanistic pathways by which microbial cues influence oogenesis are distinct from those activated by dietary perturbations. Unlike high-sugar diets, which impair GSC proliferation and disrupt ISC-to-enterocyte differentiation via insulin dysregulation and oxidative stress pathways^51^, microbial cues promote GSC proliferation and remodel gut cellular composition, including increased ISC activity and a shift toward EE-lineage progenitors (Fig. 3), potentially enhancing endocrine signaling to the ovary and brain (Fig. 2A, B, Fig. 4E,F, Fig. 5). This interpretation aligns with previous reports that microbial conditions influence serotonin production in an EE cell–dependent manner^75^. Our scRNA-seq analyses further support this distinction: under M+ conditions, insulin pathway activity was attenuated, as indicated by *foxo* downregulation, whereas the juvenile hormone-binding protein *Jhbp5* was upregulated, consistent with enhanced endocrine output rather than insulin-driven growth (Fig. 7H,I and Supplementary Fig. 7B,H). Together, these findings indicate that microbial cues engage metabolic and hormonal pathways that are distinct from those activated by dietary sugar, enabling coordinated and tissue-specific effects on stem cell proliferation and differentiation.

Our findings reveal that microbial modulation of gut metabolism contributes to the coordination of circadian programs across the gut–brain axis, thereby promoting germline stem cell activity and supporting oogenesis. Under M+ conditions, multiple core clock genes, including *tim*, *per*, and *vri*, were upregulated in both the gut and brain (Fig. 4C,D). Functional perturbation of these genes in the gut and brain suppressed the microbe-induced GSC increase (Fig. 4A,B). In addition, gut-specific knockdown of glycolytic and PPP enzymes reduced circadian gene expression in the head (Fig. 4E,F), indicating that peripheral metabolic activity can influence central circadian regulation. Previous studies have established mechanistic links between circadian clock regulation and oogenesis^76^ as well as a bidirectional influence of circadian regulation and metabolism. Circadian oscillations are tightly linked to redox and metabolic cycles, including PPP-derived NADPH rhythms across multiple biological systems^55,77^, and glycolysis and PPP can directly influence circadian oscillations through redox- and metabolite-sensitive mechanisms^39,42,43,56,78–80^. In addition to these intrinsic metabolic–circadian interactions, microbial metabolites, such as short chain fatty acids, amino acids, and polyamines, have been shown to interact with host metabolic and circadian pathways in a time- and diet-dependent manner via the gut in both *Drosophila* and mice, thereby contributing to temporal regulation of host metabolism^54,81–84^. Whether these microbiome-dependent metabolic and circadian effects contribute directly to reproductive regulation has remained largely unexplored.

Our findings further suggest that microbial modulation of gut metabolism influences circadian regulation across the gut–brain axis, which contributes to GSC increase (Fig. 4A,B,E,F). Overall, these outcomes support an additional, metabolism-linked upstream layer of regulation mediated by the gut in reproduction. Nevertheless, the precise hierarchy among these signaling pathways will require further clarification, including whether circadian regulation functions independently or cooperatively with metabolic pathways in controlling reproductive physiology.

At the level of ovarian output, endocrine signaling in ovarian somatic cells represents a key step linking systemic cues to niche activity. To understand how systemically generated signals alter activity in the ovary, we leveraged single-cell transcriptomic analyses to resolve cell- and stage-specific responses to microbial cues. Previous studies have shown that germline metabolic requirements and stress responses are dynamically reprogrammed across differentiation, enabling selective engagement of pathways such as glycolysis or the PPP in a stage-dependent manner^50,67^. In this context, microbial cues appear to be integrated through these pre-existing metabolic states, resulting in differential and stage-specific transcriptional outputs rather than uniform activation across the lineage (Fig. 7C-G; Supplementary Fig. 7D). In contrast, somatic niche lineages, including FSCs, ECs, and TF/CCs, displayed stable transcriptional profiles across conditions, consistent with their established role in localized signaling rather than extensive transcriptional remodeling^85–90^. Nonetheless, module scoring revealed subtle pathway-level shifts, particularly in ECs that directly interface with the early germline (Supplementary Fig. 7G), indicating fine-tuned metabolic and hormonal adjustments.

Collectively, our findings establish a mechanistic model (Fig. 7J) in which environmental microbes coordinate reproductive physiology through inter-organ communication. The metabolic shift induced by microbial colonization alters circadian clock gene expression along the gut–brain axis while also potentiating ecdysone and JH signaling in ovarian somatic cells. Future studies will be required to identify the specific microbial metabolites that initiate these systemic responses and to determine whether transcriptional changes in clock genes are accompanied by alterations in circadian behavior or physiology. Together, our findings reveal how environmental microbes shape reproductive physiology by linking metabolic state, circadian gene expression, and endocrine signaling across tissues.

## Materials and methods

### Drosophila strains

All fly stocks were maintained at 25 °C on standard cornmeal food^21^. The CS9515 strain was used as the wild-type (WT). All fly strains used in this study are listed in materialsData1.

### *Drosophila* microbe-sensitization assays

Microbe-sensitization assay are performed as previously described^21^. To generate microbe-rich (M+) conditions, five male pupae were placed into fresh food vials. After 3-4 days, the males were removed and replaced with five female pupae. For microbe-poor (M-) conditions, no male pupae were introduced into the vials. Female pupae were cultured for three days prior to the evaluation of oogenesis. For each experiment, three M+ and three M− vials were prepared. Germline stem cell (GSC) numbers were analyzed using a chi-square test (\*\*\*\**P* ≤ 0.001, \*\*\**P* ≤ 0.005, \*\**P* ≤ 0.01, \**P* ≤ 0.05, n.s., nonsignificant (*P* > 0.05)).

### RNA-seq analysis in brain, gut and ovaries

Brains, guts, and ovaries were dissected from females maintained under M+ or M− conditions for 3 days. Total RNA was extracted as described above and treated with DNase at 37 °C for 10 min. RNA quality was assessed using a Bioanalyzer, and all samples showed RNA integrity numbers (RINs) suitable for sequencing (8.4-10.0). RNA-seq was performed by GENEWIZ (AZENDA) following the standard Illumina sequencing protocol.

### Analysis for RNA-seq data

Raw 150 bp-paired-end RNA-seq reads in FASTQ format were quality-checked and trimmed using fastp (ver. 0.20.1) with default parameters unless otherwise specified. Adapter sequences were automatically detected, and low-quality bases were trimmed from both read ends. Quality-trimmed paired-end reads were aligned to the *Drosophila melanogaster* reference genome (BDGP ver.6.32, FlyBase) using STAR (ver. 2.5.2b) with the default parameters unless otherwise specified.

Genome index was pre-built from the reference FASTA files and the corresponding gene annotations (GTF) files. The resulting BAM files were sorted and indexed using SAMtools (ver. 1.10) and used for downstream quantification with featureCounts (ver.2.0.0). Gene-level read counts were generated using the featureCounts from the Subread package in paired-end mode, with exon features and gene-level summarization.

The resulting count matrix was used for differential expression analysis with DESeq2 (ver. 1.44.0) in R (ver. 4.4.0)^91^. Raw counts were combined with sample metadata (tissue type and experimental condition) and converted into a DESeqDataSet object using DESeqDataSetFromMatrix. Counts were normalized and variance-stabilized using the variance stabilizing transformation (VST).

Differential expression analysis was performed using a negative binomial generalized linear model including condition and batch effects in the design formula. Differential expression between the M+ and M− conditions was assessed using Wald tests. Log₂ fold changes were shrunk with the *apeglm* method to improve effect size estimation for lowly expressed genes. *P* values were adjusted for multiple testing using the Benjamini–Hochberg procedure, and genes with an adjusted *P* value < 0.05 were considered significant.

### Visualization of RNA-seq data

PCA was conducted on the 500 most variable genes across all samples to visualize global transcriptional differences according to experimental factors, including tissue type and condition (M+ and M−), using the plotPCA function in DESeq2. The percentage of variance explained by the first two principal components is indicated on the axes of the resulting PCA plots. Differential expression results were visualized using volcano plots, with log₂ fold change plotted on the x-axis and −log₁₀(*P* value) plotted on the y-axis. Vertical dashed lines indicate the fold-change threshold (log₂FC = 1.5). Selected genes of interest were highlighted in red.

### Quantitative real-time PCR (qRT-PCR)

Total RNA was extracted from gut, brain, or head tissues using TRIzol (Thermo Fisher Scientific). Reverse transcription was performed using 1 μg of total RNA using SuperScript III reverse transcriptase (Thermo Fisher Scientific). qRT-PCR was conducted on a StepOnePlus Real-Time PCR system (Thermo Fisher Scientific) with at least three biological replicates. Gene expression levels were quantified using the ΔΔCT method^92^, and fold-change values were calculated relative to *rp49* or actin. Primer sequences are listed in the materialsData2.

### Immunostaining

Ovaries were dissected and fixed in 5.3% paraformaldehyde (PFA) in PBS for 10 min, followed by three 20-min washes in 0.2% Triton X-100 in PBS (PBX). Samples were then blocked in 4% BSA with 0.2% PBX for 40 min and incubated overnight at 4 °C with primary antibodies (materialsData1 for details). After three 15-min washes in PBX, Alexa Fluor-conjugated secondary antibodies diluted in 0.4% BSA in PBX were applied overnight at 4 °C. Following three additional 15-min washes in PBX, nuclei were stained with DAPI (1:500 in wash buffer), and samples were mounted in Fluoro-KEEPER mounting medium (Nacalai Tesque). Gut immunostaining was performed using the same procedure. Images were acquired using a Zeiss LSM900 or LSM780 confocal microscope with 63× or 40× objectives and processed using Zeiss microscope software (ZEN blue). Germline stem cell (GSC) counts were obtained using an Olympus Axiovert microscope. Statistical significance of GSC numbers was evaluated using a chi-square test (****P ≤ 0.001, ***P ≤ 0.005, **P ≤ 0.01, n.s., not significant, P > 0.05).

### Counting colony forming units (CFUs) from donor or recipient flies

To quantify microbial load in whole flies based on colony-forming unit (CFU) counts, we followed established protocols^93^ with minor modifications, as described previously^21^. Flies were homogenized in 125 μL of PBS using sterile pestles for 1 min, and the homogenate was diluted to a final volume of 1 mL. 10 μL of serial dilutions (1:9, 1:81, and 1:729) were applied to the mannitol and MRS plates. Plates were Incubated at 30 °C for 2 days, and the numbers of colonies were subsequently counted.

### Mass spectrometry and data processing

Mass spectrometry–based metabolomic analysis was performed on dissected female guts (10 mg per sample) from M+ and M−conditions (n = 5). Anionic and cationic metabolites were quantified using IC-MS and LC-MS, respectively. Metabolite abundances were normalized to total metabolite content per sample. Replicate consistency was assessed by Pearson correlation analysis, and outlier samples were excluded. Differential metabolite analysis was conducted using Welch’s t-test, and fold changes were calculated as log₂(M+/M−) and visualized using heatmaps and box plots. Pathway-level enrichment of PPP, glycolysis, and TCA cycle metabolites was evaluated using gene set enrichment analysis. Detailed procedures are described in the Supplementary methods.

### Quantification of receptor activation

Female flies of the genotypes *hs-GAL4-EcR; UAS-nlacZ* ^57,58^ or *hs-GAL4-Gce; UAS-nlacZ* female flies^21^ with or without *zw* RNAi or *Mal-B2* RNAi driven by *TKg-Gal4* or *mex-1-Gal4,* were cultured for 3 days under M+ or M-conditions. Flies were then heat-shocked at 37 °C for 1 h in a block incubator and allowed them to recover overnight at 25 °C. Animals were dissected and stained with SPiDER-βGal to visualize and quantify fluorescence derived from LacZ activity. The enzymatic activity of LacZ converts SPiDER-βgal into an reactive intermediate that covalently binds to the surrounding proteins through nucleophilic interaction, generating a fluorescent signal. Fluorescence intensity by SPiDER-βGal was normalized to the corresponding DAPI signal. Images were acquired using a Zeiss LSM780 confocal microscope with 63× or 40× objectives and processed using Zeiss microscope software (ZEN blue). Statistical significance was assessed using a Wilcoxon rank sum test (\*\*\**P* ≤ 0.005, \**P* ≤ 0.05, n.s, nonsignificant (*P* > 0.05)).

### Single-cell RNA-seq sample preparation

Single-cell RNA-seq libraries were prepared from ovaries dissected from 60 Canton-S females maintained under M+ or M− conditions for 3 days (N = 2). Non-vitellogenic ovaries (up to stage 9) were dissociated based on a published protocol with minor modifications^60^. Dissociated cell suspensions from each experimental condition were pooled and adjusted to a concentration of approximately 1,000 cells/μL prior to loading. Libraries were prepared using the Chromium Single Cell 3′ Library & Gel Bead Kit v3 (10x Genomics) according to the manufacturer’s instructions and sequenced on an Illumina NovaSeq 6000 platform using the recommended paired-end configuration (28 bp for Read 1, 8 bp for the index read, and 90 bp for Read 2). Detailed procedures are described in the Supplementary methods.

### Single-cell RNA-seq data analysis and visualization

Single-cell RNA-seq data were processed using the Cell Ranger pipeline (ver. 7.1.0; 10x Genomics) to generate gene–cell UMI count matrices aligned to the *Drosophila* melanogaster reference genome. Downstream analyses were performed in R (ver. 4.4.0) using Seurat (ver. 5.3.0^94^). Cells with low transcript complexity (<100 genes), potential doublets (>6,000 genes), or elevated mitochondrial transcript content (>5%) were excluded using Cell Ranger and Seurat, resulting in approximately 7600∼8000 high-quality cells from M+ samples and 5000∼7000 cells from M- samples per dataset (SupplementaryTable2). Data integration, dimensionality reduction, and clustering were performed using standard Seurat workflows. PCA and UMAP were used for dimensionality reduction, and clustering was performed using the Louvain algorithm at a resolution of 1.5. Cluster identities were assigned based on established marker genes for *Drosophila* ovarian cell types^60,61,63,64^.

Cell-type identities were assigned based on established marker genes validated in previous ovarian single-cell studies^60,61,63,64^. These include germline cells spanning early differentiation stages, escort cells (ECs), follicle stem cells (FSCs), pre-follicle cells, polar and stalk lineages, as well as stage-specific follicle-cell populations. Germline clusters were identified by expression of canonical markers such as *vasa*^95–97^, whereas somatic populations were distinguished by lineage-and stage-specific transcriptional signatures representative, including *Wnt4* and *br*^60,61,98,99^. Additional markers enabled confident annotation of polar and stalk lineages, hemocytes, and muscle cells^60–62^. Cell-type annotations were validated by heatmaps of marker gene expression as well as dot plots, feature plots, and violin plots and were consistent across biological replicates and experimental conditions.

Differential expression analysis was conducted for each annotated cell type using the DESeq2 framework implemented in Seurat to compare M+ and M− conditions. Genes were considered differentially expressed if they showed an adjusted P value < 0.05 and an absolute log₂ fold change > 0.1. Pathway-level changes were assessed using predefined gene sets for metabolic and signaling pathways, and module score–based analyses and UCell analyses were used to evaluate pathway activity in a cell-type-specific manner. Functional enrichment analyses were performed as a complementary analysis and are described in the Supplementary methods. Detailed data processing and analysis procedures are also provided in the Supplementary methods.

## Acknowledgments

We thank the Advanced Genome Support (AGS) program of the Platform for Advanced Genome Science Research (PAGS), supported by the Ministry of Education, Culture, Sports, Science and Technology (MEXT), Japan and research funds from the Yamagata Prefectural Government and Tsuruoka City. We also thank Dr. Masaya Matsui for ovary dissections for single-cell analysis and Dr. Kazumi Abe for technical support with the 10x Genomics Chromium platform. We acknowledge the Bloomington *Drosophila* Stock Center, The Vienna *Drosophila* Resource Center and the Kyoto Stock Center for providing fly stocks. We are grateful to Dr. Ritsuko Morita for discussions on the single-cell RNA-seq analysis workflow.

## Author contributions

Conceptualization, R.S., J.Y.Y. and T.K.; Methodology, R.S., A.H., K.A., H.M.,Y.S, S.K. and J.Y.Y.; Investigation, R.S. Z.M., R.M., H.M. and K.A.; Writing, R.S.; Review & Editing, R.S., S.K., J.Y.Y and T.K.; Funding Acquisition, R.S., A.K., J.Y.Y. and T.K.

## Data and material availability

NGS data sets have been deposited to the DNA Data Bank of Japan (DDBJ). BioProject Accession: PRJDB40088 and PRJDB12675. All fly strains and antibodies generated for this study are available upon request.

